# Accurately modelling RNase H-mediated antisense oligonucleotide efficacy

**DOI:** 10.1101/2025.10.29.685292

**Authors:** Barney Hill, Maisie R. Jaques, Remya R. Nair, Nicola Whiffin, Matthew J. A. Wood, Stephan J. Sanders, Peter L. Oliver, Alyssa C. Hill, Carlo Rinaldi, the UPNAT Consortium

## Abstract

Antisense oligonucleotides (ASOs) are a powerful class of drugs with the potential to treat a wide range of human diseases. However, the prediction of ASO efficacy remains challenging, as large-scale and costly experimental screens are typically required to identify optimal candidates for a specific therapeutic target. To address this challenge, we compiled ASO Atlas, a database comprising 188,521 RNase H-mediated ASO sequences targeting 334 unique genes with corresponding knockdown efficacy measurements extracted from published patents. Using ASO Atlas, we trained OligoAI, a deep learning model capable of jointly modelling RNA target context, ASO sequence, sugar and backbone chemistries, and dosage to predict *in vitro* efficacy. We experimentally validated OligoAI by targeting KCNT2, achieving a 5.72-fold reduction in screening effort compared to random selection. ASO Atlas provides the first systematic resource to rigorously evaluate hypotheses regarding key parameters in ASO design, including sequence composition, chemical modifications, and target region selection. Both ASO Atlas and OligoAI have been made freely accessible through an online web-tool with the aim of facilitating the accelerated optimisation of ASO design.

## Introduction

Antisense oligonucleotides (ASOs) are short, synthetic, single-stranded nucleic acids designed to selectively hybridise with target RNA sequences and modulate gene expression. ASOs can reduce target RNA levels or alter the splicing patterns of target RNA transcripts. The former is achieved by recruitment of RNase H, an enzyme that cleaves the RNA strand of the DNA-RNA heteroduplex formed between the ASO and its RNA target^2,3^. RNase H-dependent ASOs typically employ a ‘gapmer’ design ^35^. This design features a central gap of 8-10 DNA nucleotides, which are essential for RNase H recognition and cleavage, flanked by wings containing 3-5 nucleotides modified at the 2’ position of ribose (e.g., 2’-*O*-methoxyethyl, 2’-MOE; constrained ethyl, cEt), which increase binding affinity and stability^4^ and enhance the ASO’s drug-like properties^3^. The backbone is typically uniformly modified with the phosphorothioate (PS) linkage, which replaces one non-bridging oxygen atom of the natural phosphodiester (PO) linkage with a sulfur atom to confer nuclease resistance on the ASO. This strategy effectively balances the requirement for RNase H activity with the need for enhanced affinity and stability. Gapmer structures are often denoted by the lengths of the 5’ wing, DNA gap, and 3’ wing (e.g., 5-10-5).

Initially proposed as therapeutics over four decades ago^5^, ASOs utilising gapmer designs have recently gained significant clinical momentum, with landmark approvals such as volanesorsen for Familial Chylomicronemia Syndrome and inotersen for Transthyretin Amyloidosis, demonstrating their viability for treating rare genetic disorders^6,7^. As their safety profile becomes increasingly established, opportunities for treating ultra-rare genetic diseases continue to expand^8–11^.

Despite these clinical successes, the determinants of ASO efficacy remain poorly understood. Effective ASO design requires optimising two key components: the pharmacophore, which is defined by the nucleotide sequence and dictates target specificity, and the dianophore, which includes the pattern of chemical modifications, which influence the drug-like properties^4^. However, predicting the combined impact of sequence and chemistry on efficacy remains a significant challenge. With gene transcripts containing thousands of potential ASO target sites, and there being many potential combinations of oligonucleotide length and chemical composition, current pre-clinical development strategies are heavily burdened by costly experimental screening efforts.

Current computational predictive models fail to capture the complex structure-activity relationships of ASOs^12^, and these approaches are further hampered by the lack of standardised, large-scale training datasets. Furthermore, existing computational tools are often constrained by proprietary data that cannot be independently evaluated. In this study, we present ASO Atlas, an extensive resource compiled from ASO screening data reported throughout the patent literature. Leveraging this comprehensive dataset, we have developed OligoAI, a machine-learning approach to jointly model how ASO sequence and chemical modifications influence *in vitro* efficacy, demonstrating a data-driven approach that enhances the identification of potent ASO candidates.

## Results

### ASO Atlas: A large-scale dataset of gapmer ASOs with corresponding in vitro efficacy measurements

The ASO Atlas comprises a diverse collection of 188,521 RNase H-mediated gapmer ASO sequences extracted from 417 United States Patent and Trademark Office (USPTO) patents published between 2001 and 2025. To construct this dataset, we implemented a multi-stage computational pipeline that began by manually labelling tables containing ASO knockdown efficacy data from patent applications made by IONIS Pharmaceuticals. We then employed a two-step Large Language Model approach: OpenAI’s gpt-5 converted raw XML patent tables into standardised CSV format using Python scripts, followed by systematic annotation of ASO sequences, chemical modifications, target genes, and quantitative efficacy measurements based on our dataset schema requirements with SQL. This dataset contains nucleotide sequence composition, chemical modifications, target gene, and corresponding quantitative reverse transcription polymerase chain reaction (qRT-PCR) RNA abundance measurements across multiple cell lines. Manual review of 100 randomly selected entries across 100 patents revealed no numerical transcription errors, and only minimal errors in chemical modification transcriptions (3% contained at least one error). The dataset features experiments conducted in a diverse range of cell types including A-431 (*N* = 44,855), HepG2 (*N* = 23,428) and SH-SY5Y (*N* = 18,192) and features 334 unique gene targets. The gene targets with the most ASOs were *F12* (*N* = 5,874), *DGAT2* (*N* = 5,326), and *PMP22* (*N* = 5,273) (Figure 1c). Among ASOs with chemistry annotations, 52.6% were 5-10-5 2’-MOE gapmers and 30.4% were 3-10-3 cEt gapmers (with varying PS backbone patterns), while the remaining proportion comprised diverse designs including alternative gapmer lengths (e.g., 4-10-4, 5-9-5) and mixed MOE/cEt chemistries. While the *atlas* spans from 2001 to 2025, we observed a notable increase in the volume of patented sequences in recent years (see 2018-2024, Figure 1b). The dataset exhibits a broad distribution of efficacy values (Figure 1d) with a median knockdown efficacy of 45.0%. Importantly, 17.4% of ASOs show <10% knockdown, indicating substantial inclusion of ineffective sequences. This representation across the full efficacy spectrum suggests that patent filings document comprehensive screening results, rather than selectively reporting successful candidates, thus providing a robust training signal for machine learning models.

**Figure 1.**
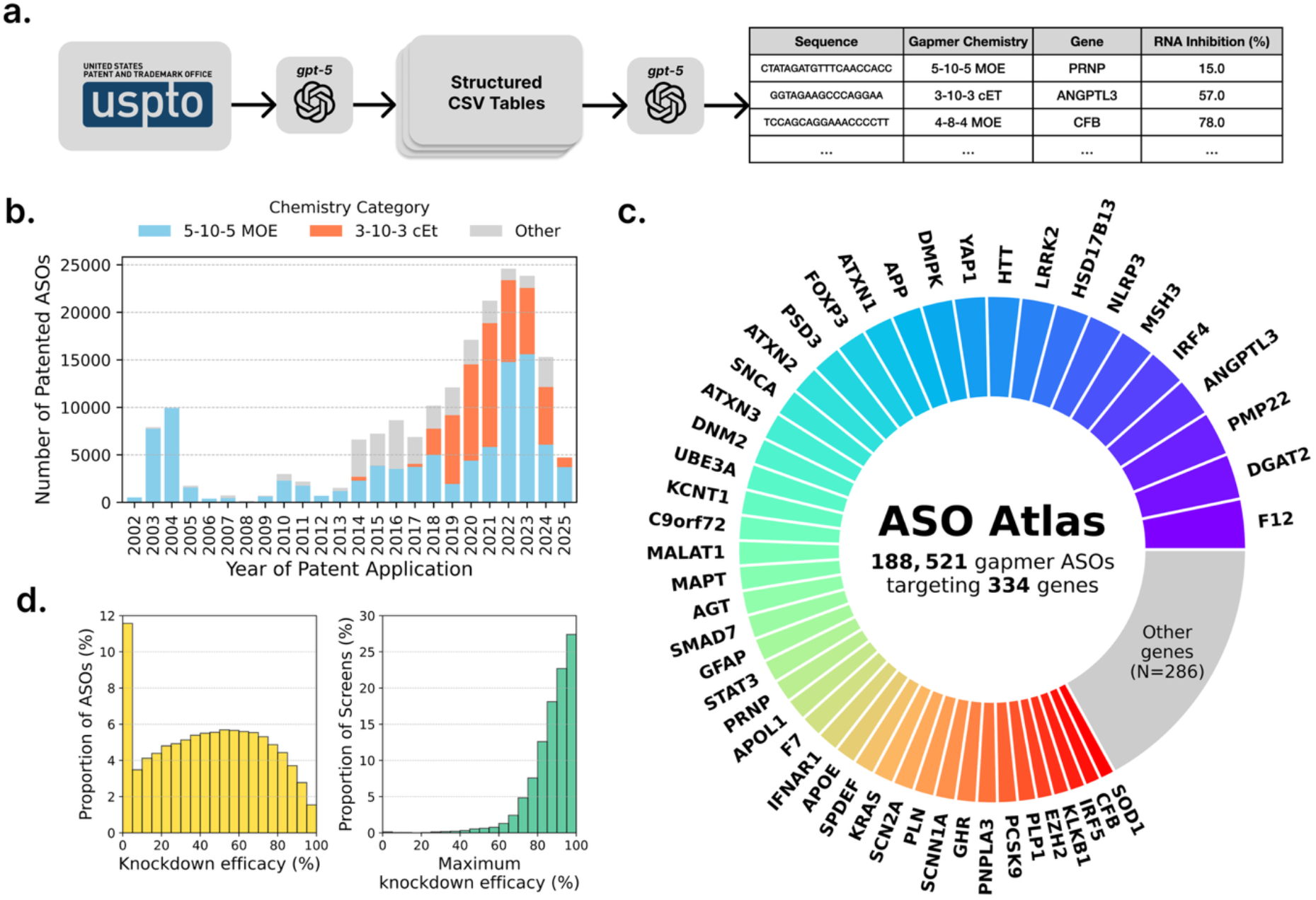
Construction and composition of the ASO Atlas dataset. (a) Pipeline for creating the ASO Atlas dataset, showing extraction of ASO data from USPTO patents using gpt-5 for initial structured table extraction, followed by annotation of sequence, chemistry, target gene, and knockdown measurements. (b) Temporal distribution of patented ASO sequences from 2001-2025, showing a peak in patented sequences in 2023. Blue represents 5-10-5 MOE modified ASOs, orange represents 3-10-3 cEt ASOs and grey represents other chemistries. (c) Donut chart of ASO distribution across genes, demonstrating concentrated development efforts on select therapeutic targets. Major genes with ≥1,500 ASOs are labeled individually, while the ‘Other genes’ category encompasses 286 genes with < 1,500 ASOs. (d) RNA knockdown distributions for all ASOs (left, yellow) showing a broad efficacy range including ineffective ASOs, and maximum knockdown per screen (right, green), indicating that a large proportion of screens achieve high knockdown with at least one ASO.

### Target site characteristics influence gapmer ASO efficacy

We systematically evaluated multiple factors influencing ASO efficacy, including nucleotide sequence composition, genomic location, regulatory element overlap, and RNA secondary structure. To identify position-specific nucleotide patterns associated with efficacy, we performed multivariate linear regression analysis for each nucleotide at each position along the ASO sequence, separately for 5-10-5 MOE (all PS) and 3-10-3 cEt (all PS) chemistries. After Bonferroni correction, we identified 83 significant position-specific nucleotide effects that varied across chemistry types (Figure 2a,b). For example, cytosines showed opposite effects between chemistries: 13/20 positions were significantly positively associated with knockdown efficacy in 5-10-5 MOE ASOs, while 12/16 positions showed negative association in 3-10-3 cEt ASOs. The only pattern shared across both chemistry types was thymine showing positive association at positions 7-12 (5-10-5 MOE) and 1-9 (3-10-3 cEt) suggesting chemistry-dependent modulation of sequence composition effects.

**Figure 2.**
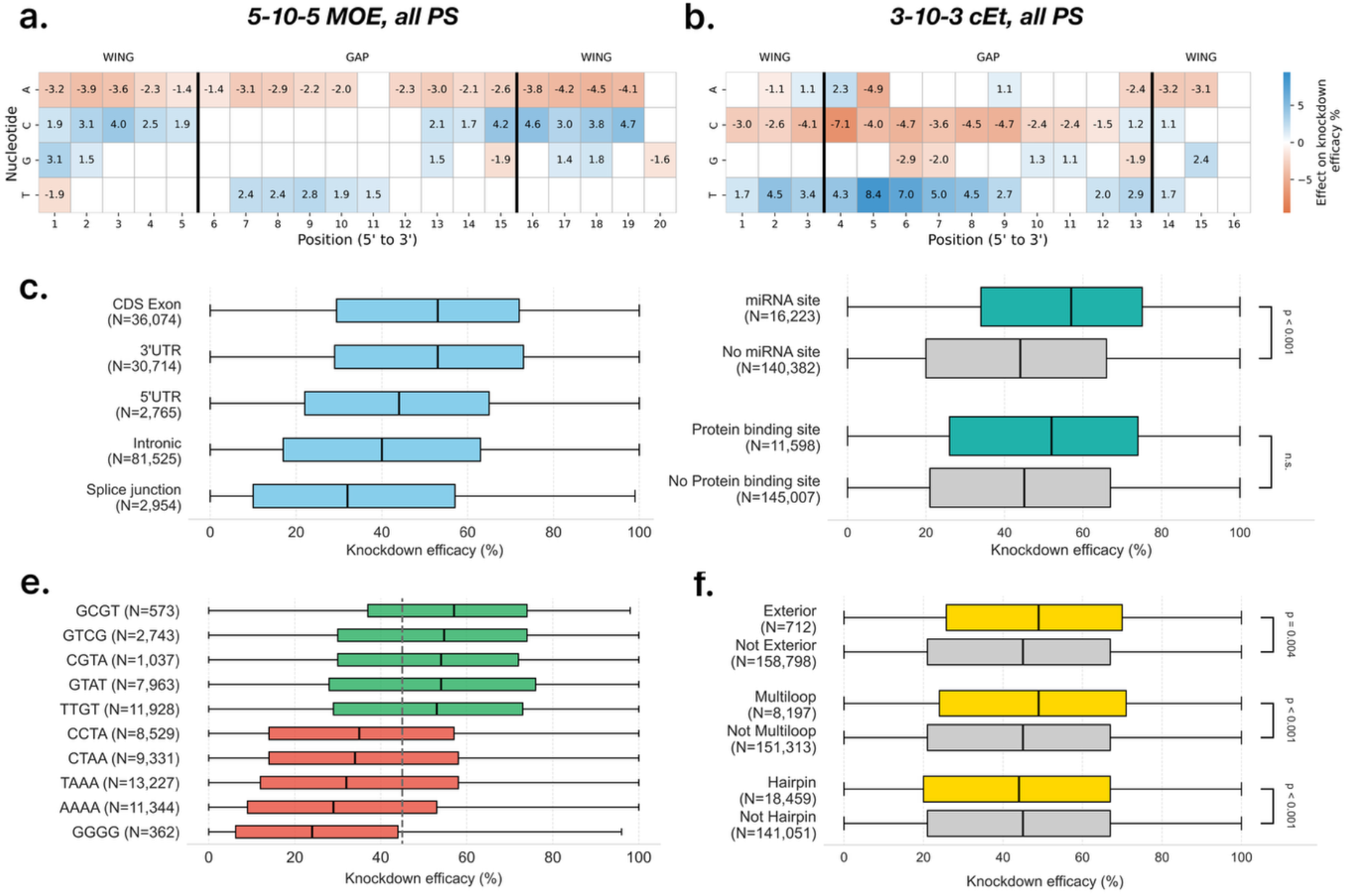
Analysis of ASO knockdown efficacy across genomic regions and sequence contexts. (a,b) Heatmaps of position-specific nucleotide effects on ASO efficacy for 5-10-5 MOE, all PS and 3-10-3 cEt, all PS chemistries, respectively. Effects from linear regression are shown only where significant after Bonferroni correction. Colors indicate effect magnitude on knockdown efficacy percentage, with blue representing positive effects and orange representing negative effects on efficacy. (c) Comparison of efficacy distributions between ASOs overlapping different genomic regions. Regions are presented in order of median efficacy, and all group differences are significant (Mann-Whitney U, see Methods). (d) Comparison of efficacy distributions between ASOs overlapping miRNA binding sites as reported by TarBase and protein binding sites as reported by RBP-Tar^16,17^. (e) Bar chart displaying top/bottom-5 median efficacy differences of 4-mer motifs (All significant after accounting for screen confounding with linear model, see Methods), with green bars showing positive effect motifs and red bars showing negative effect motifs. (f) Comparison of efficacy distributions between ASOs overlapping RNA secondary structure elements including multiloop, external, hairpin as annotated by ViennaRNA 2.0 using ± 150 bp context around the ASO target^*14*^.

Analysis of ASO efficacy across different genomic target region classifications revealed statistically significant variation in target knockdown (*P* < 0.001, Kruskal-Wallis test). We observed that ASOs targeting splice junction regions exhibited the lowest median knockdown efficacy (32.0% [IQR: 10.0%–57.0%]), followed by intron-targeting ASOs (40.0% [IQR: 17.0%– 63.0%]) and 5’UTR-targeting ASOs (44.0% [IQR: 22.0%–65.0%]). In contrast, both 3’UTR and exon-targeting ASOs demonstrated the highest median knockdowns (53.0% [IQR: 28.0%– 73.0%] and 53.0% [IQR: 29.0%–72.0%] respectively). We note that higher efficacy is observed for loci present in both mature RNA transcripts and pre-mRNA (coding exons, 3’UTR, and 5’UTR) compared to those only present in pre-mRNA (splice sites and introns), see Figure 2a.

To further investigate the mechanisms underlying these regional differences, we first examined overlaps with microRNA (miRNA) and RNA-binding protein sites, given their enrichment in untranslated regions where we observed the varying efficacy^13^. After accounting for regional variation in ASO efficacy with a multivariate linear regression analysis controlling for genomic region type, only miRNA binding site overlap (*β* = 4.43%, *P* < 0.001) was identified as a significant predictor of efficacy.

Next, we investigated the role of RNA secondary structure on efficacy based on overlap with ViennaRNA-predicted pre-mRNA secondary structure elements (using ±150 bp context)^14^. After adjusting for screen-to-screen variability (using a linear mixed model), ASOs targeting unpaired regions showed 8.5% higher knockdown efficacy compared to those targeting base-paired regions, indicating that target accessibility significantly influences ASO performance (*P* < 0.001). Predicted structure types showed distinct effects on efficacy. Hairpin structures were associated with decreased median knockdown efficacy (43.0% vs 45.0%, *N* = 18,459 vs 141,051, *P* < 0.001, Mann-Whitney U test), but multiloop structures enhanced performance (49.0% vs 45.0%, *N* = 8,197 vs 153,313, *P* < 0.001, Mann-Whitney U test). Exterior loop structures also improved efficacy (49.0% vs 45.0%, *N* = 712 vs 158,798, *P* = 0.0036, Mann-Whitney U test) (Figure 2d).

Finally, to identify sequence motifs that influence ASO knockdown efficacy, we analysed all possible 4-mer nucleotide patterns while statistically controlling for variability between experimental screens. The ‘GGGG’ motif showed the most pronounced significant association; ASOs containing this sequence exhibited a markedly lower median knockdown efficacy (24.0%), a finding consistent with previous studies^15^ (Figure 2c). Several other motifs, including ‘AAAA’ (29.0%), ‘TAAA’ (32.0%), ‘CTAA’ (34.0%), and ‘CCTA’ (35.0%), were also significantly associated with lower efficacy (all p < 0.001). In contrast, motifs such as ‘TTGT’ (53.0%), ‘GTAT’ (54.0%), ‘CGTA’ (54.0%), ‘GTCG’ (54.7%), and ‘GCGT’ (57.0%) were significantly associated with higher efficacy.

### OligoAI: A deep learning model for ASO efficacy prediction

To leverage the sequence, structural, and positional determinants identified in ASO Atlas, we developed OligoAI (https://sitlabs.org/OligoAI), a transformer-based deep learning model that jointly encodes ASO nucleotide sequence, target RNA context, chemical modifications, dosage and transfection method to predict *in vitro* target RNA knockdown (Figure 3a)^18^. The model takes the ASO sequence and its target pre-mRNA context (±50 nucleotides flanking the hybridisation site), position-specific chemical modifications, including 2’-MOE and cEt sugar modifications, as well as PS and PO backbone linkages, and experimental parameters such as dosage and transfection method.

**Figure 3.**
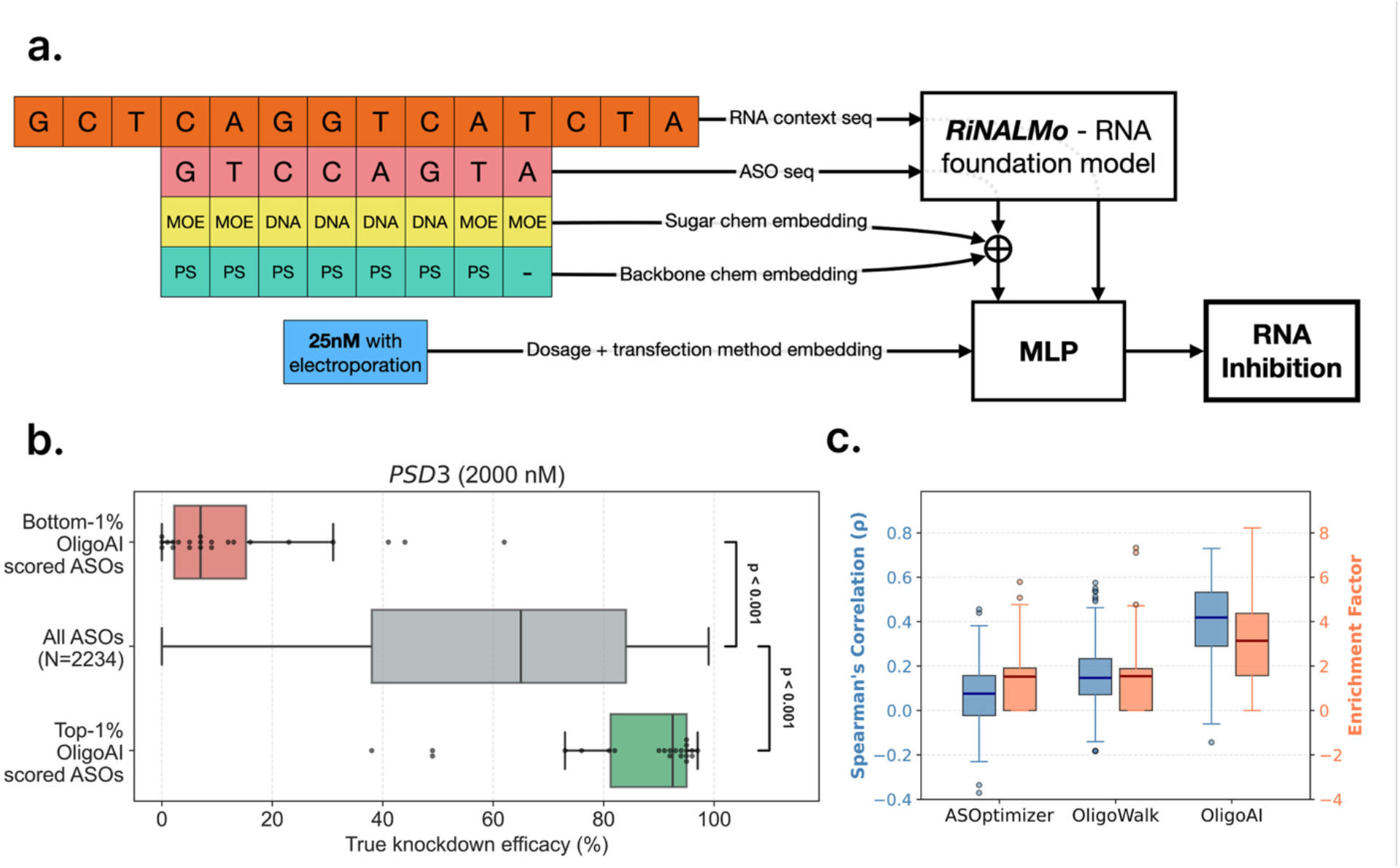
OligoAI model architecture and performance evaluation. (a) Architecture of OligoAI showing the integration of multiple feature modalities. ASO sequences and their target RNA context (±50 nucleotides) are encoded using the pre-trained RiNALMo-giga transformer model. Position-specific sugar modifications and backbone chemistry are encoded through learned embeddings and concatenated with RiNALMo sequence representations. The final prediction is generated by a MLP that integrates the pooled ASO representation, pooled context representation, and dosage-scaled transfection method information to predict percent knockdown. (b) Validation on a large held-out screen (PSD3: *N* = 2,234) showing that OligoAI-predicted top 1% ASOs achieve significantly higher knockdown than screen average, while predicted bottom 1% ASOs show significantly lower knockdown (One-Sample Wilcoxon Test). (c) Distribution of Spearman correlation (*ρ*) and enrichment factor across *N* = 299 held-out test screens, comparing OligoAI performance to baseline methods (ASOptimizer, OligoWalk). Enrichment factor is the ratio of hit rate in the predicted top 10% to the baseline hit rate of 10%, where hits are defined as ASOs in the true top 10% by measured inhibition.

The architecture integrates these feature modalities through a multi-stage encoding process. ASO sequences and their target RNA contexts are independently encoded using RiNALMo-giga, a 650-million parameter bidirectional transformer pre-trained on 36 million non-coding RNA sequences, to capture local secondary structure accessibility and sequence context effects. Chemical modifications are represented through learned embeddings and concatenated with the RiNALMo-encoded ASO sequence representations. These combined features are processed through a bottleneck network to learn cross-modal sequence-chemistry interactions. The resulting ASO and context representations are then globally pooled and integrated with dosage-scaled transfection method embeddings through a multi-layer perceptron to generate final efficacy predictions.

When evaluated across 299 held-out test screens, OligoAI achieved a median Spearman correlation of 0.419 [IQR: 0.290–0.533] between predicted and measured knockdown efficacy values, with a 3.14× enrichment factor for identifying high-performing ASOs (meaning the top 10% of OligoAI predictions contained 3.14 times more ASOs that were truly in the top 10% by measured knockdown efficacy compared to random selection) (Table 1). This substantially outperforms previous state-of-the-art thermodynamics-based models ASOptimizer (*ρ* = 0.076, 1.53× enrichment) and OligoWalk (*ρ* = 0.147, 1.55× enrichment), which rely primarily on hybridisation energetics and target site accessibility^12,19^

**Table 1:** Model performance comparison (median [IQR] values reported across validation test set, N=299 qRT-PCR screens)

| Model | Spearman $\rho$ | Enrichment factor |
| --- | --- | --- |
| ASOptimizer | 0.076 (-0.023–0.158) | 1.53× (0.00–1.92) |
| OligoWalk | 0.147 (0.071–0.234) | 1.55× (0.00–1.89) |
| **OligoAI** | **0.419 (0.290–0.533)** | **3.14× (1.57–4.38)** |

To assess OligoAI’s practical utility for identifying hits for *in vitro* gene screens with many possible sequence designs, we evaluated its performance on a large held-out screen targeting PSD3 (*N* = 2,234 ASOs). The model demonstrated robust discriminative ability: while the overall screen achieved a median knockdown efficacy of 65.0%, OligoAI’s top 1% scored ASOs (*N* = 22) achieved a median knockdown efficacy of 92.5%, (*P* < 0.001, one-sample Wilcoxon test) (Figure 3b), highlighting OligoAI’s potential to substantially reduce experimental screening burden in therapeutic ASO development.

### Experimental validation of OligoAI prioritisation in KCNT2 screening

To test the utility of OligoAI in a real-world experimental screening setting, we generated a virtual library of 200,374 20-mer unique gapmer ASOs targeting the Potassium Sodium-Activated Channel Subfamily T Member 2 (*KCNT2*), a gene associated with a rare developmental epileptic encephalopathy^20,21^.

To confirm that OligoAI prioritises the previously identified biologically relevant features, we examined whether the top 1% ranked *KCNT2* ASOs were enriched for genomic and sequence characteristics associated with efficacy. Top-ranked candidates showed significant enrichment for exonic targets (*N* = 118/2,006 in top 1% vs *N* = 5,510/198,368 in remaining sequences; fold enrichment = 2.12, *P* = 1.43 × 10^−13^, Fisher’s exact test), presence of top-5 sequence motifs (TTGT, GTAT, CGTA, GTCG, GCGT; *N* = 570/2,006 vs *N* = 32,707/198,368; fold enrichment = 1.72, *P* = 2.61 × 10^−40^), and miRNA binding site overlap (*N* = 22/2,006 vs *N* = 658/198,368; fold enrichment = 3.31, *P* = 2.35 × 10^−6^). Finally, proportion of unbound target RNA structure showed little difference between groups (mean = 0.423 vs 0.421; fold change = 1.00, *P* = 0.048, Mann–Whitney *U* test, structure predicted using ViennaRNA 2.0 using ± 150 bp context around the ASO target^14^).

Two groups of ASOs were defined for *in vitro* testing: 32 randomly sampled from the entire library and 18 randomly sampled from the 1% of ASOs with the highest OligoAI scores. To determine a suitable experimental dose for the ASO testing, a pilot experiment using two randomly chosen ‘active’ ASOs was carried out by transfection in HeLa cells. After testing a range of doses and incubation times, 30 nM dosing for 48-hours was selected as it provided the most suitable dynamic range of *KCNT2* knockdown by the ‘active’ ASOs with no likely floor effects (Supplementary Fig. 1). In the subsequent complete library screening experiment, the relative *KCNT2* knockdown level was determined for all 50 ASOs in parallel, providing a ranking of *in vitro* efficacy (Supplementary Fig. 2). ASOs ranked in the top 1% by OligoAI achieved superior target knockdown efficacy (median = 81% [IQR: 64–88%]) compared to randomly selected ASOs (median = 36% [IQR: 11–62%]) (Mann-Whitney U test, *P*=1.1×10^−4^). These results demonstrate OligoAI’s ability to effectively identify high-performing ASO candidates (Figure 4). Follow-up dose-response studies confirmed these results were reproducible, with IC50s in the low nanomolar range (Supplementary Fig. 3).

**Figure 4.**
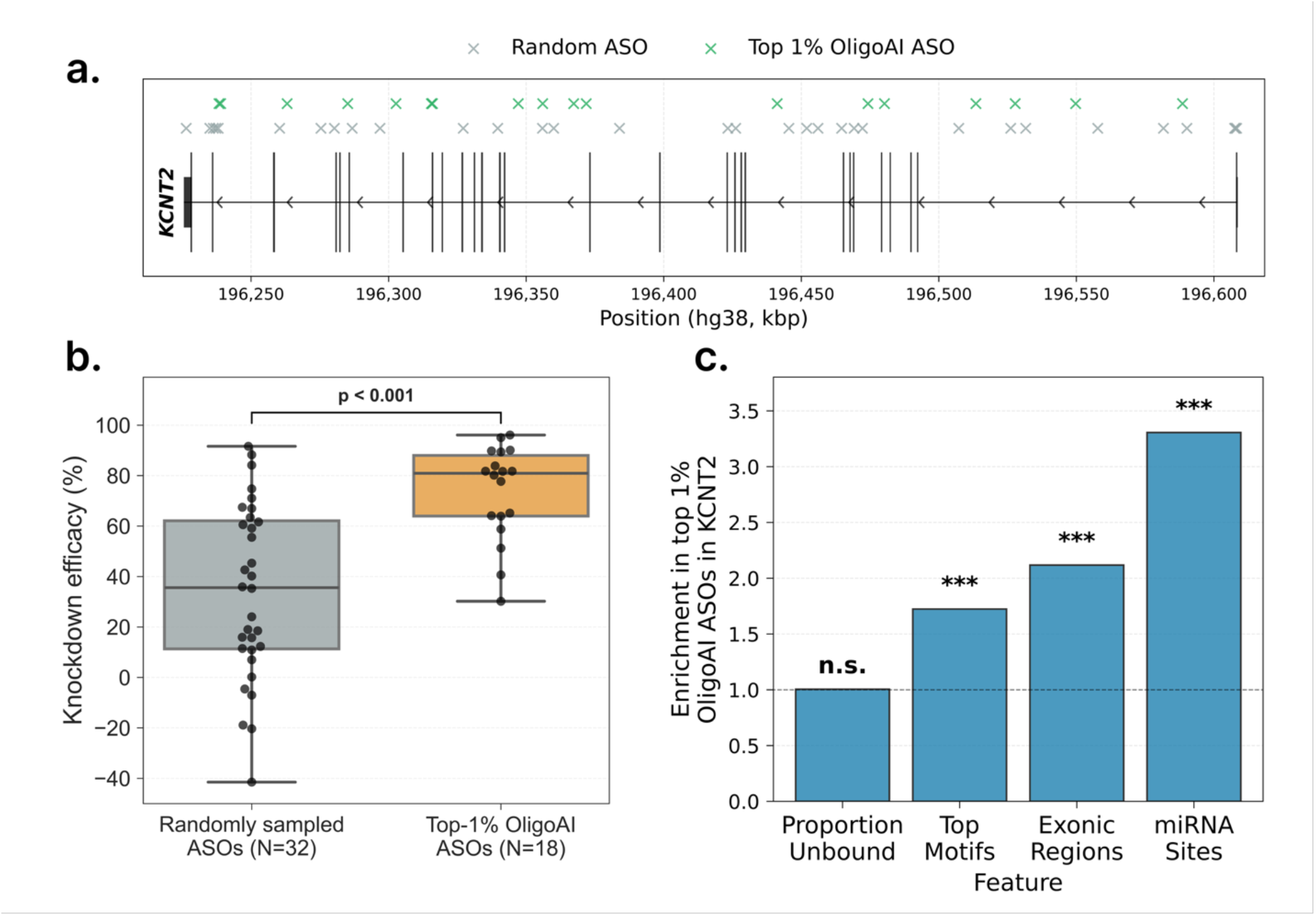
Experimental validation of OligoAI prioritisation in *KCNT2* screening. Schematic representation of the *KCNT2* transcript structure showing the genomic locations of ASOs sampled from two groups: randomly selected ASOs (*N* = 32), and top 1% OligoAI-ranked ASOs (*N* = 18). (b) Comparison of *KCNT2* knockdown efficacy across the ASO groups, expressed as relative target gene expression following transfection in HeLa cells. Top 1% OligoAI-ranked ASOs demonstrated significantly superior performance compared to randomly selected ASOs (Mann-Whitney U test, *P* = 1.1×10^−4^), with median relative expression levels of 0.19 versus 0.64, respectively. (c) Barplot showing the enrichment of biologically relevant features (exonic targeting, top-5 motifs, miRNA binding sites, and unbound RNA structure) within the top 1% ranked *KCNT2* ASOs compared to remaining sequences. Asterisks indicate statistical significance after Bonferroni correction (***, *p* < 0.001; n.s., not significant).

To quantify the practical screening efficiency gains, we performed bootstrap analysis (*N* = 10,000 iterations) to estimate the screening effort required for random selection to achieve equivalent performance to OligoAI’s top candidates. Our analysis revealed that to match the median knockdown efficacy of our 18 top-scored ASOs, a standard random screening approach would require testing 103 ASOs (95% CI: 53–179). This represents a 5.72-fold reduction in required screening effort (95% CI: 2.94–9.94-fold), demonstrating substantial cost savings and accelerated candidate identification for ASO development workflows.

## Discussion

Our systematic analysis of 188,521 gapmer ASOs reveals key principles governing RNase H-mediated ASO efficacy and establishes a predictive framework for therapeutic design. The observed variation in efficacy across genomic regions, combined with secondary structure analysis, points to RNA accessibility as a central determinant of ASO performance. The reduced efficacy observed for hairpin-targeting ASOs compared to those targeting multiloop and exterior loop regions suggests that stable secondary structures present thermodynamic barriers to ASO hybridisation. This interpretation is further supported by the positive association between miRNA binding site overlap and ASO efficacy, since miRNA target sites must be structurally accessible to enable miRNA binding, suggesting that ASOs similarly benefit from targeting accessible regions^22^. Together, these findings indicate that successful ASO design must consider not only sequence complementarity but also the structural and genomic context of the target site.

ASO design has historically relied on heuristic trial-and-error approaches with limited datasets, with computational tools primarily focusing on thermodynamic parameters while largely ignoring the chemical modifications that define modern therapeutics^23,24^. Existing algorithms such as OligoWalk predict hybridisation stability based on nearest-neighbour thermodynamics but fail to account for the phosphorothioate backbones and sugar modifications (2’-MOE, cEt) that comprise the majority of clinical ASO candidates^19^. More recently, ASOptimizer has emerged as a two-component framework that addresses both sequence selection and chemical modification optimization separately^12^. Its sequence engineering module employs a linear regression model incorporating three thermodynamic features: target site binding free energy, off-target binding potential, and secondary structure accessibility. ASOptimizer also includes a separate chemistry optimization module based on graph neural networks that optimizes modification patterns given a fixed sequence. However, this bifurcated approach has notable limitations. Our systematic analysis revealed strong interactive effects between sequence composition and chemistry type: position-specific nucleotide associations with efficacy were almost entirely different between 5-10-5 MOE and 3-10-3 cEt chemistries (Fig 2a,b), with only thymine showing consistent patterns across both. These chemistry-specific sequence preferences demonstrate that nucleotide composition and chemical modifications do not act independently but rather likely interact to determine ASO efficacy. By treating sequence selection and chemistry optimization as independent problems, ASOptimizer cannot capture these complex, non-linear interactions between nucleotide sequence, chemical modifications, and target RNA structure that govern ASO efficacy. In our comparative evaluation across 299 held-out screens, ASOptimizer’s sequence module achieved only modest predictive performance (Spearman ρ=0.076), substantially lower than OligoAI (ρ=0.419). OligoAI’s improved performance likely stems from several key architectural differences including jointly modelling sequence and chemistry rather than treating them as independent optimization problems and leveraging a pre-trained RNA language model (RiNALMo) which has been shown to perform across various RNA modelling tasks.

While OligoAI represents a significant advance in ASO design prediction, several important limitations highlight areas for future development. First, the current model is limited to predicting RNase H-dependent ASOs. For many targets, this modality presents a large sequence design space, providing both greater opportunity for model training and a stronger need for computational optimization to guide experimental screening efforts. Additionally, our reliance on automatic extraction of patent-derived data represents a constraint, with LLM-based annotation introducing chemistry annotation errors in 4% of cases. This is a tolerable rate for machine learning approaches but with scope for future improvement. Finally, future extensions to the OligoAI platform will incorporate in vivo efficacy data, off-target effects, and toxicity assessments, all critical factors that ultimately determine therapeutic success.

In summary, we have created the ASO Atlas, the largest publicly available database of ASOs with experimentally validated efficacy data, comprising 188,521 gapmers targeting 334 genes. Through systematic analysis of this resource, we generated critical insights into the sequence, positional, and chemical modification features that govern ASO performance *in vitro*, revealing that exonic targets demonstrate superior efficacy. We translated these insights into OligoAI, a deep learning prediction tool validated through holdout screens and experimental KCNT2 testing, achieving a 5.72-fold reduction in experimental screening burden. By making both the ASO Atlas database and OligoAI prediction tool freely available to the research community, we provide resources that will substantially reduce the cost and time associated with ASO development, accelerating the translation of antisense therapeutics from target identification to clinical application. When combined with emerging experimental platforms such as patient-derived organoid screening systems^25^, this work enables more rapid progression toward personalised ASO therapeutics, ultimately enhancing the accessibility of precision medicine approaches for patients with rare genetic disorders.

## Methods

### Dataset collection and processing

#### Patent selection and initial data extraction

We constructed ASO Atlas by mining the USPTO Bulk Data Storage System (BDSS) for ASO efficacy data from 2001-2025. After filtering for patents published by either ISIS or IONIS Pharmaceuticals, we manually annotated tables containing knockdown efficacy data. From these tables, we also included the preceding five paragraphs as contextual information.

#### Table structure normalisation and validation

We developed a multi-stage natural language processing pipeline to convert unstructured XML patent tables into standardised formats. The full prompts provided to each model are detailed in Supplementary Section, and the final dataset conforms to a structured schema, which is fully defined in Supplementary Table 3. For table structure recognition, we employed OpenAI’s gpt-526 with verbosity=‘low’ and reasoning=‘low’ to create a Python script to convert the raw XML table into normalised CSV format^26^. Subsequently, we performed schema validation using gpt-5 to generate SQL commands that concatenate the ASO sequence information and quantitative efficacy measurements (either inhibition percentage or UTC percentage) across tables.

#### Chemistry annotation and sequence characterisation

For each validated entry, we implemented comprehensive annotation across three main stages. First, we used gpt-5 to systematically extract and standardise chemical modification patterns from patent text through chemistry annotation. Our annotation framework captured three types of chemical modifications (phosphorothioate backbone, 2’-MOE sugar, and cEt sugar), as well as unmodified backbone (PO) and fully unmodified DNA sequences. For each ASO, we recorded both modification types and their precise positions along the sequence, with the model being provided with the five previous text paragraphs and row-specific data for this task. Manual review of 100 randomly selected entries revealed a 3% error rate in chemical modification transcriptions, with errors including the exclusion of mC modifications and a 4-9-4 gapmer being incorrectly recorded as a 5-7-5. These errors were manually corrected prior to downstream analysis. Second, we performed target information extraction using gpt-5 to annotate entries with metadata including target RNA name, HGNC gene symbol, experimental cell line, ASO dosage (nM), cell density, and transfection method extracted from the five previous text paragraphs. Finally, we conducted genomic target mapping by mapping each ASO sequence to its target genomic location using the Ensembl 110 human genome assembly^27^. For each target gene, we selected the Ensembl canonical transcript, and ASO target sites were identified by searching for reverse complementary sequence matches between the ASO sequence and the pre-mRNA transcript.

#### Dataset quality assurance

We implemented rigorous quality control measures throughout the curation process. For sequence filtering, we restricted our dataset to ASOs with sequence lengths between 12-30 nucleotides and removed tables with duplicate sequence-inhibition pairs to prevent data redundancy. To ensure extraction accuracy, we performed manual verification by cross-referencing 100 randomly selected entries across 100 different patents. No errors were detected in ASO sequences and inhibition values in our validation sample, with a 97% accuracy rate in chemistry annotation. Regarding model selection rationale, we employed gpt-5 for annotation tasks based on benchmarking that demonstrated superior task accuracy and cost-effectiveness compared to alternative models such as gpt-5-mini. The final curated dataset comprises 188,521 ASOs from 417 patents, with comprehensive annotation of nucleotide sequences, chemical modifications, target genes, and experimental conditions (Table 2).

**Table 2:** Antisense oligonucleotide modifications present in ASO Atlas.

| Modification Type | ASO Count | Unique Patents |
| --- | --- | --- |
| PS Backbone | 185,585 | 409 |
| 2'-MOE | 126,620 | 391 |
| cEt | 77,516 | 53 |
| Any | **188,521** | **417** |

#### Position-wise nucleotide association testing

For each position along the ASO sequence and each nucleotide (A, C, G, T), we constructed a linear regression model with knockdown percentage as the dependent variable. The independent variables included: (1) a binary indicator for the presence of the specific nucleotide at the given position, (2) target gene identity, (3) transfection method, and (4) dosage (nM). This approach controlled for confounding effects of experimental conditions while isolating the contribution of each position-nucleotide combination to ASO efficacy.

Position-nucleotide combinations with insufficient variation (fewer than 5 ASOs containing or lacking the nucleotide at that position) were excluded from analysis. For each regression model, we extracted the coefficient and *P*-value corresponding to the nucleotide indicator variable. To account for multiple hypothesis testing across all position-nucleotide combinations, we applied Bonferroni correction, with statistical significance defined as corrected *P* < 0.05.

#### Comparison of knockdown efficacy by genomic target region

ASO target regions were classified hierarchically into mutually exclusive categories with the following priority: 5’UTR, 3’UTR, splice junction (overlapping an exon-intron boundary), CDS exonic and intronic as defined by Ensembl 110 canonical transcripts. ASOs overlapping miRNA or protein binding sites were defined by at least 10 nucleotides shared between the ASO target region and experimentally validated binding sites to reduce false positives. miRNA binding site regions were sourced from TarBase v9 using PAR-CLIP or HITS-CLIP data^16^, while protein binding sites were obtained from RBP-Tar using eCLIP data (*P* < 0.05, Bonferroni-corrected for 515,644 sites)^17^.

Comparisons of efficacy across different genomic target regions (5’ UTR, Splice Junction, Intron, Exon, 3’ UTR) utilised the Kruskal-Wallis H-test for an overall assessment of differences among the five region types. Following a statistically significant result, post-hoc pairwise comparisons between specific region types were performed using Dunn’s test, with p-values adjusted for multiple comparisons using the Bonferroni correction method. All statistical analyses were performed using Python (v3.10) with the SciPy library (v1.15.1)^28^ for the Kruskal-Wallis and Mann-Whitney U tests.

#### Motif Analysis

To identify ASO motifs associated with knockdown efficacy while controlling for experimental batch effects, we performed a systematic analysis of all 256 possible 4-mer motifs. For each motif, a multiple linear regression model was fitted to predict knockdown percentage based on the motif’s presence, with the experimental screen (custom_id) included as a categorical covariate to account for screen-to-screen variability. To ensure adequate statistical power, only motifs present in at least 15 sequences were analysed. We applied a strict Bonferroni correction for multiple comparisons, establishing a significance threshold of *p* < 0.05/256 (≈ 1.95 × 10^−4^). The direction of a motif’s effect (positive or negative) was determined from its regression coefficient, and all motifs presented in this study were significant under these criteria.

### Computational models for ASO efficacy prediction

#### Deep learning model architecture

We implemented OligoAI, a transformer-based architecture that leverages the pre-trained RNA language model RiNALMo to predict ASO efficacy. The model processes multiple input modalities to capture both sequence and experimental context:

- **Sequence representations**: ASO sequences and their target RNA context (ASO binding site ±50 flanking nucleotides) are encoded using RiNALMo-giga, a pre-trained bidirectional transformer model specialised for RNA sequences. Representations are extracted from the final transformer layer, yielding 1,280-dimensional embeddings per nucleotide position. The ±50 nucleotide context window was selected to capture local RNA secondary structure elements that determine ASO binding site accessibility, consistent with established approaches for local structure prediction^32,34^, while maintaining computational tractability for the transformer architecture.
- **Chemistry track**: Position-specific sugar modifications (MOE, cEt, DNA) are encoded through learned embeddings (16 dimensions).
- **Backbone track**: Phosphorothioate (PS) or phosphodiester (PO) linkages are represented through separate embeddings (8 dimensions).
- **Experimental conditions**: Transfection method (electroporation, gymnosis, lipofection, other) is encoded through learned 4-dimensional embeddings and element-wise multiplied by log-transformed dosage (log1p) to capture method-specific dose-response relationships.

The architecture integrates these features through a multi-stage process. First, ASO sequence representations from RiNALMo (1,280-dim) are concatenated with chemistry (16-dim) and backbone embeddings (8-dim)^18^. These combined features (1,304-dim total) pass through a bottleneck network (linear 1,304-dim to 128, ReLU, dropout p=0.2, linear 128-dim to 1,280, ReLU) to learn cross-modal representations^29^. Both ASO and context representations are then processed through global pooling layers that project to 64 dimensions while handling variable-length sequences via masked mean pooling. The final prediction is made by a 3-layer MLP with 128 hidden dimensions that combines the pooled ASO representation (64-dim), pooled context representation (64-dim), and method-scaled dosage embedding (4-dim) through fully connected layers with ReLU activations and dropout (p=0.3).

#### Training and evaluation methodology

The model was trained to directly predict percent knockdown values using a mean squared error (MSE) regression objective. Target knockdown values were standardised (zero mean, unit variance) using a StandardScaler fitted on the training set, with predictions inverse-transformed to the original scale for evaluation. We initialised the model with pre-trained RiNALMo-giga weights and employed a gradual unfreezing schedule during fine-tuning to preserve learned RNA representations while adapting to the ASO efficacy prediction task. At epoch 0, only randomly initialised components were unfrozen (prediction head, chemistry embedder, backbone embedder, transfection method embedder). At epoch 3, we unfroze the pre-trained RiNALMo final layer normalisation and transformer blocks 6-39. OligoAI was optimised using the Adam optimizer with a learning rate of 5 × 10^−5^, weight decay of 0, and a linear learning rate schedule decaying to 5 × 10^−6^ (end factor = 0.1) over the total training steps^30^. Training proceeded for 10 epochs with a batch size of 64 samples, employing gradient clipping (max norm = 0.5) to ensure stable convergence. The model was trained with mixed precision (16-bit) on NVIDIA L40S GPUs (48GB memory) to improve computational efficiency. The final model was selected based on the epoch achieving the lowest validation set MSE.

We evaluated the model using an 80/10/10 train/validation/test split, with stratification performed at the patent level to prevent data leakage between related ASO sequences. This patent-level splitting ensures that all ASOs from a given patent appear in only one split, addressing the tendency for patents to contain structurally and functionally related oligonucleotides. Model performance was assessed using multiple metrics: coefficient of determination (*R*^2^), mean absolute error (MAE), root mean squared error (RMSE), and Spearman rank correlation (*ρ*) calculated per experimental screen and averaged across screens. Enrichment factor was defined as the ratio of the hit rate in the predicted top 10% to the baseline hit rate of 10%, where hits were defined as ASOs in the true top 10% by measured knockdown.

#### Model comparisons

To compare the relative performance of OligoAI we used ASOptimizer and OligoWalk^12 19^. Specifically, from ASOptimizer we implemented the linear regression model from the sequence engineering module with the published regression parameters: *ŷ* = *a*_0_ + *a*_1_*x*_1_ + *a*_2_*x*_2_ + *a*_3_*x*_3_, where *a*_0_ = −1.077, *a*_1_ = −0.037, *a*_2_ = 0.019, *a*_3_ = 1.422, *x*_1_ represents the target Gibbs free energy change calculated by miRanda^31^, *x*_2_ represents the mean of the top-10 off-target Gibbs free energy calculated against canonical protein coding pre-mRNA transcripts by miRanda^31^, and *x*_3_ represents the secondary structure accessibility (ratio of unhybridised nucleotides) calculated by mFold^32^. From OligoWalk we used the oligowalk_overall metric, which incorporates multiple thermodynamic terms including duplex stability (Δ*G*_*duplex*_), target mRNA secondary structure stability (Δ*G*_*target_structure*_), oligo self-structure (Δ*G*_*intra_oligomer*_), and the oligo-oligo dimer (Δ*G*_*inter_oligomer*_) to predict overall binding affinity.

### Cell culture, ASO treatment and gene expression analysis

HeLa cells were grown on 96-well plates (Thermo Scientific, 167008) at a seeding density of 10,000 cells per well in 100 *μ*l media (DMEM with GlutaMAX (Gibco, 61965026), 10% FBS (Gibco, A3840402) and 1% Penicillin Streptomycin (Gibco, 15070063)). The following day, cells were treated with individual ASOs at the doses indicated. The media was replaced with 100 *μ*l fresh media before dosing and 20 *μ*l of transfection mix (0.15 *μ*l Lipofectamine RNAiMAX (Thermo Scientific, 13778150) with the ASO dose in OptiMEM (Gibco, 31985062) media). For the initial quantitative (q)RT-PCR assay optimisation, ASOs were dosed for 24, 48, or 72 hours at 3, 10, 30 or 100 nM. For the ASO library screen, ASOs were dosed for 48 hours at 30 nM. Three biological replicate wells were used for each treatment. Controls included untransfected cells (OptiMEM), RNAiMAX transfection with no ASO and a non-targeting control with a matching 5-10-5 2’-MOE gapmer design containing PS bonds throughout and 5’methyl-C / dC bases (5’-AGTCGCACACGTCTATACGC-3’). For dose-response analysis, a log10 dosing scale was used from 0.1 nM to 1 *μ*M for 48 hours as above. ASOs were synthesised by ATDBio or Integrated DNA Technologies (IDT) with QC by HPLC and LC / MS.

After ASO dosing, total RNA was extracted using the MagMAX mirVana Total RNA Isolation Kit (Applied Biosystems, A27828). RNA concentration was measured using the Qubit RNA High Sensitivity Assay Kit (Invitrogen, Q32855) and cDNA synthesis was carried out using the High-Capacity cDNA Reverse Transcription Kit (Applied Biosystems, 4368813). Quantitative RT-PCR was performed using Fast SYBR Green Master Mix (Applied Biosystems, 4385618) on a CFX Opus 384 Real-Time PCR System (Bio-Rad). The following cycling conditions were used: enzyme activation at 95°C for 20 seconds, followed by 40 cycles of denaturation at 95°C for 3 seconds, annealing and extension at 60°C for 30 seconds. The qRT-PCR primer sequences were as follows: 5’-GTGCAGACACTCTTCAGGTTG-3’ and 5’-AGCCTCTCTCCCGTTCTTTC-3’ for *KCNT2*; 5’-AGTTCTGTGGCCATATGCTTAGTAG-3’ and 5’-AAACAACAATCCGCCCAAAGG-3’ for the reference gene *HPRT*. qRT-PCR data were analysed using the comparative Ct (ΔΔ*Ct*) method: the Ct values of the target gene were normalised to the Ct values of the reference gene. Relative gene expression levels were calculated using the formula 2^−ΔΔ*Ct*^. All reactions were performed in technical triplicates per biological replicate and the average *Ct* values used for analysis.

## Data and Code Availability

The ASO Atlas dataset scripts and OligoAI training code are available at https://github.com/barneyhill/aso_atlas and https://github.com/barneyhill/OligoAI. The trained model can be downloaded at https://huggingface.co/barneyhill/OligoAI. An interactive web interface for ASO design is available at https://sitlabs.org/OligoAI.

