## supplementary_material for "Accurately modelling RNase H-mediated antisense oligonucleotide efficacy"

***ASO Atlas Dataset Schematic.*** Description of the fields, data types, and definitions for each entry in the ASO Atlas dataset.

| Field Name | Data Type | Description |
|---|---|---|
| aso_sequence_5_to_3 | String | The nucleotide sequence of the ASO. |
| inhibition_percent | Float | The measured percentage of target RNA inhibition. Typically 0-100 although there are some values < 0 where upregulation was observed. |
| chemistry | Object | A object containing a list of all sugar and backbone modifications |
| custom_id | String | A string refering to the location of the referenced patent table file. |
| target_mrna | String | The name used to refer to the target mRNA. |
| target_gene | String | The HUGO gene name corresponding the the target_mrna |
| cell_line | String | The cell-line used for the screen. |
| dosage | Float | The dosage of the administered ASO in nM. |
| cells_per_well | Integer | The number of cells per well used in the screen. |
| transfection_method | String | The method of introducing the ASO into the cells. |

**LLM Prompts for Data Extraction**

To ensure the reproducibility of our data extraction pipeline, this section details the verbatim prompts used for table normalisation with gpt-5.

```
Prompt 1

## Task
Write a Python 3.11 script to convert OCR-extracted table XML data into a
structured CSV format.

## Output Format
Return a Script object with:
- `pyscript`: Complete Python conversion script as a string

## Function Requirements
- The Python script should contain a function `xml_to_csv(xml_str: str) ->
str`
- You do need to return if __name__ ... w/ example usage - just our
function.

## Technical Requirements
### Dependencies
- Uses only Python standard library + `re` module
```

```
### Column Name Rules
- Preserve meaning of original column headers in new csv. i.e "UTC
Untreated control group (%)" to "utc_untreated_control_group_pct" /
"Inhibition (%)" to "inhibition_pct"
- Make SQL-compatible: underscores for spaces, no dots, lowercase
- Hardcode column names (no need to dynamically generate)

### Data Handling
- Use "NA" for missing/empty cells
- Properly escape CSV values (quotes, commas, newlines)
- Use comma delimiter
- Sometimes   is used in the XML, this should be replaced with a
space in the CSV.

## Domain Context
- UTC = "Untreated Control" percentage - this is not inhibition
- Preserve scientific notation and decimal precision
- Don't interpret abbreviations unless obvious

## Quality control
- To ensure correct rows let's strip newspace and capitalise the sequence
column. It should have >=8 ATGC characters. If not skip the row.

## Preview of the input XML Structure (xml_str) - we will use the full
version as input for your script (do not return this in your output!):
Your goal is to produce a dataset of antisense-oligonucleotide sequences
and their inhibition percentages.
To do so you must stack the secondary_table with the primary_table using a
SQL command.

Required Columns in secondary_table:
1. ASO sequence (case insensitive)
2. One of:
   - inhibition/knockdown/reduction percentage
   -  UTC (Untreated Control) / RNA percentage

Transformation Rules:
- Numeric columns -> DOUBLE
- inhibition_percent =
    - Direct copy from inhibition/knockdown columns
    - 100 - UTC(%) for untreated control
- CONCAT two columns if ASO sequence is split (e.g.,sequence_part_one,
sequence_part_two)
- When using CAST be careful, some rows may not be castable to double,
hence use TRY_CAST.

Task:
1. Generate SQL:
   - Stack (INSERT INTO) secondary_table onto primary_table
   - Apply transformations as needed
   - **CRITICAL: Always use "secondary_table" as the table name in your
FROM clause**
   - **When referencing columns from secondary_table, use the exact column
```

names shown in the schema, including any special characters or numbers.**

Output Format:
- sql_command: string containing complete SQL command to stack the secondary_table onto primary_table

Data:
primary_table:
Schema:
- aso_sequence_5_to_3 (VARCHAR): 5'-3' ASO nucleotide sequence
- inhibition_percent (DOUBLE): target inhibition percentage, range 0-100

**Prompt 2**
Your goal is to produce a dataset of antisense-oligonucleotide sequences and their inhibition percentages.
To do so you must stack the secondary_table with the primary_table using a SQL command.

Required Columns in secondary_table:
1. ASO sequence (case insensitive)
2. One of:
   - inhibition/knockdown/reduction percentage
   - UTC (Untreated Control) / RNA percentage

Transformation Rules:
- Numeric columns -> DOUBLE
- inhibition_percent =
    - Direct copy from inhibition/knockdown columns
    - 100 - UTC(%) for untreated control
- CONCAT two columns if ASO sequence is split (e.g.,sequence_part_one, sequence_part_two)
- When using CAST be careful, some rows may not be castable to double, hence use TRY_CAST.

Task:
1. Generate SQL:
   - Stack (INSERT INTO) secondary_table onto primary_table
   - Apply transformations as needed
   - **CRITICAL: Always use "secondary_table" as the table name in your FROM clause**
   - **When referencing columns from secondary_table, use the exact column names shown in the schema, including any special characters or numbers.**

Output Format:
- sql_command: string containing complete SQL command to stack the secondary_table onto primary_table

Data:
primary_table:
Schema:
- aso_sequence_5_to_3 (VARCHAR): 5'-3' ASO nucleotide sequence
- inhibition_percent (DOUBLE): target inhibition percentage, range 0-100

1

**Supplementary Figures**

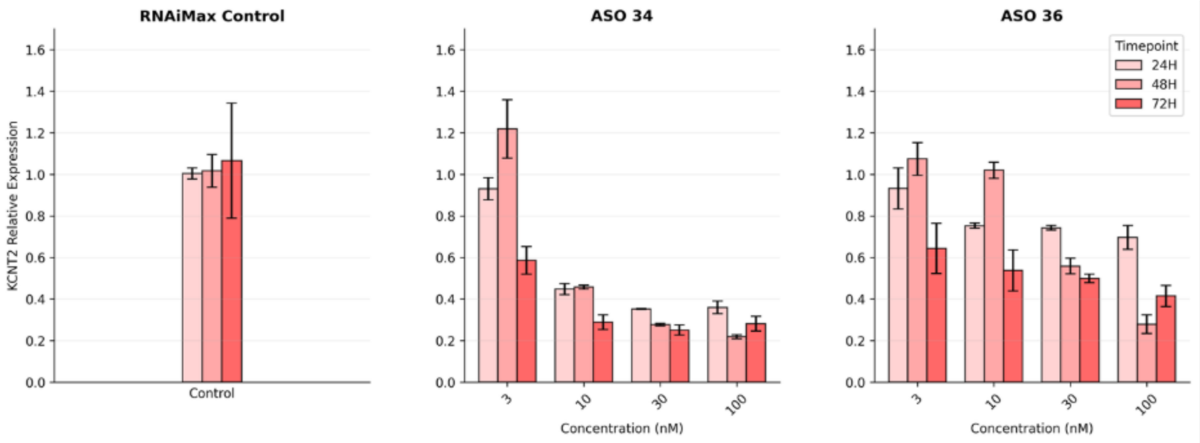


**Supplementary Figure 1. Dose- and time-dependent knockdown of *KCNT2* expression by selected ASOs in HeLa cells.** Individual panels show the relative activity for two predicted active ASOs. Cells were treated at concentrations of 3, 10, 30 and 100 nM for 24, 48 or 72 hours. *KCNT2* mRNA levels were measured by quantitative RT-PCR and normalised to *HPRT*. Data represent mean $\pm$ SEM of three biological replicates per condition. Based on these results, 30 nM treatment for 48 hours was selected as the optimal condition. Data are shown relative to transfection reagent alone (RNAiMAX).

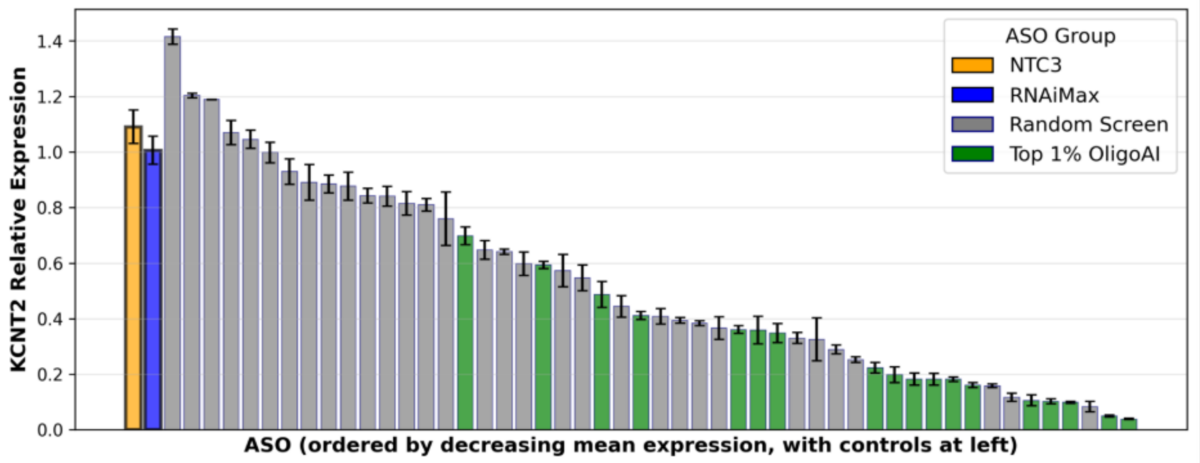


**Supplementary Figure 2.** Bar plot showing relative *KCNT2* expression levels for all 80 screened ASOs ordered by decreasing mean expression (most effective knockdown on the left). HeLa cells were transfected with individual ASOs at 30 nM for 48 hours. Relative gene expression was determined by qRT-PCR with *HPRT* as the reference gene. Error bars represent SEM from three biological replicates. Bars are colored by ASO selection group: randomly selected ASOs (grey bars, standard screen, $N$ = 32) and ASOs from the top 1% of OligoAI scores (green bars, predicted active, $N$ = 18). The distribution demonstrates that OligoAI predictions correlate with experimental efficacy, with most top 1% ASOs showing greater knockdown activity (lower relative expression) compared to bottom 1% ASOs. Data are shown relative to transfection reagent alone (RNAiMAX) and a non-targeting control (NTC) ASO of matching chemical composition.

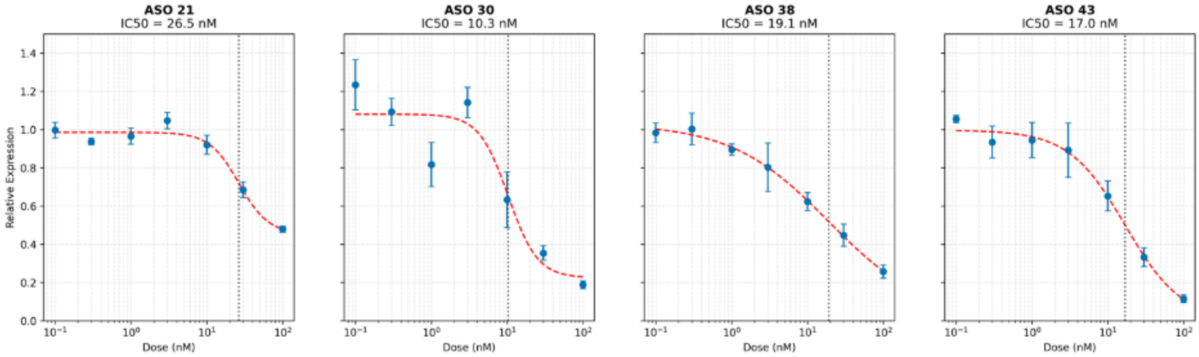


**Supplementary Figure 3. Example individual dose-response curves showing relative *KCNT2* expression levels following ASO treatment in HeLa cells.** Cells were transfected with ASOs at concentrations ranging from 0.1 nM to 1 $\mu$M and incubated for 48 hours. Each data point represents the mean $\pm$ SEM of three biological replicates. Relative gene expression was determined by qRT-PCR with *HPRT* as the reference gene. Red dashed lines show 4-parameter logistic curve fits (Hill equation). Black dotted vertical lines indicate calculated IC50 values. IC50 values demonstrate knockdown activity in the low nanomolar range.

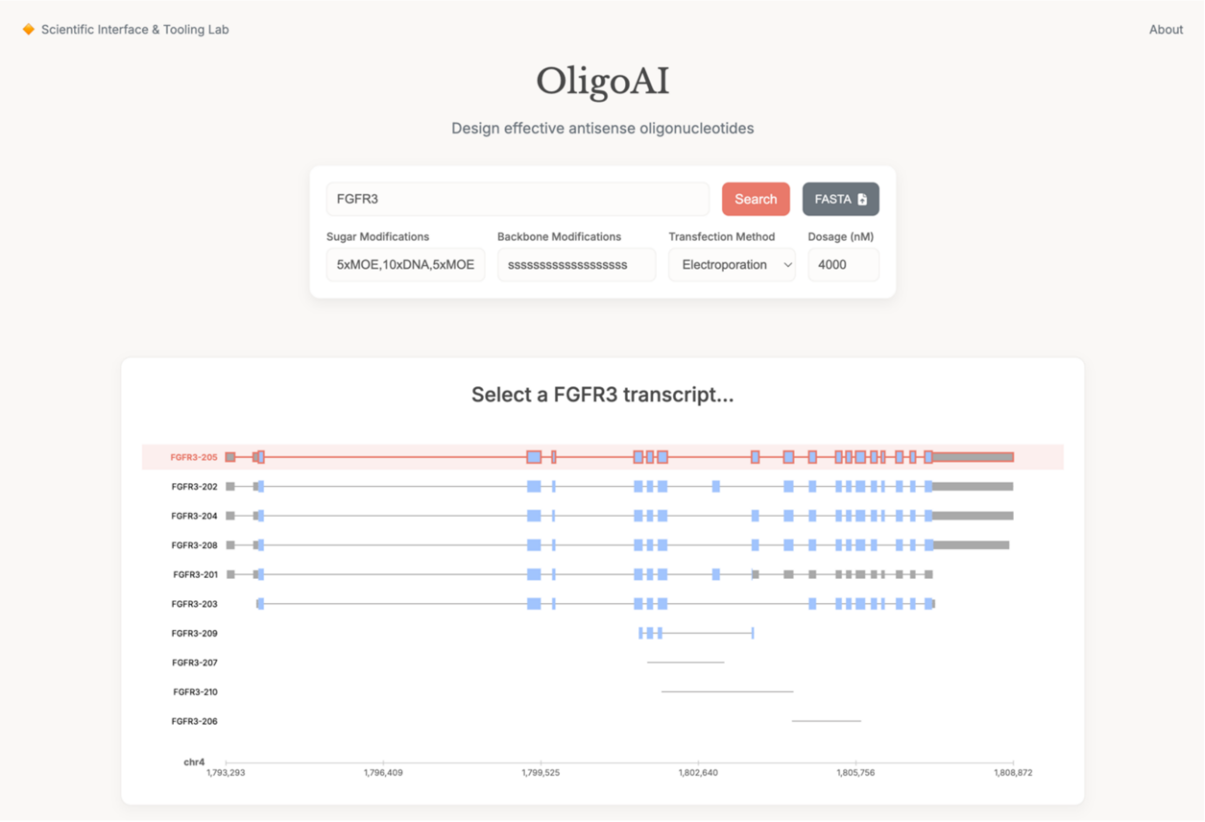


**Supplementary Figure 4. OligoAI online portal.** Users can either supply a target RNA via FASTA file upload or select an Ensembl human transcript. When a target is specified, the OligoAI model processes the target RNA using serverless GPU inference. These results are returned to the user on the same page or via an email reminder and are available to download.
